# SpatialTRACE predicts anatomical axes and regions in spatial transcriptomics and microscopy

**DOI:** 10.64898/2026.09.23.753836

**Authors:** Alexander Monell, Aishwarya A. Gogate, Dhruv Patravali, Kathleen Abadie, Gene W. Yeo, Ananda W. Goldrath, Maximilian Heeg

## Abstract

Spatial transcriptomics measures gene expression in tissue sections. However, interpretation requires anatomical maps that link gene expression and cellular composition to tissue structure. Annotating entire sections often requires extensive manual labor. We developed SpatialTRACE (Tissue Region and Axis Coordinate Estimation), consisting of graph- and image-based models, to extend annotations of a few structures into tissue-wide maps of anatomical axes and regions. SpatialTRACE-Graph combines gene-expression profiles with spatial neighborhoods to predict anatomical coordinates or region membership throughout spatial transcriptomic datasets. In held-out mouse small-intestine sections, it predicted crypt-villus and epithelial-distance axis coordinates using as few as 10 annotated training villi and identified Peyer’s patches from region annotations. SpatialTRACE-Image predicts the same anatomical axis coordinates and regions across entire tissue images from DAPI alone. This multiscale vision transformer learns from coordinate and region predictions generated by SpatialTRACE-Graph. We applied SpatialTRACE-Image to immunofluorescence images to map the anatomical distribution of antigen-specific P14 CD8 T cells responding to acute systemic infection with lymphocytic choriomeningitis virus (LCMV) Armstrong in the small intestine. Compared with the vehicle-treated section, a section treated with a retinoic acid receptor inhibitor contained fewer P14 CD8 T cells overall, with a smaller fraction in the upper-villus lamina propria and a relative enrichment in the muscularis. Overall, the SpatialTRACE models reduce repeated manual annotation and provide a route to extend anatomical maps learned from spatial transcriptomics to DAPI-containing microscopy data.

## Main

### Anatomical mapping links cellular measurements to tissue structure

Cellular states and population distributions vary with anatomical position^1,2^. For example, in the small intestine, the crypt-villus axis describes position from the crypt toward the villus tip and is associated with zonated epithelial gene-expression and metabolic programs, as well as spatial differences in stromal and immune cell populations^1–3^. Distance from the epithelium distinguishes cells near the epithelial layer from those farther into the lamina propria. Peyer’s patches are discrete lymphoid regions that support immune responses to intestinal antigens^4^. Anatomical maps can link cell phenotypes to their positions along or locations within these tissue structures.

Spatial transcriptomics measures gene expression spatially throughout a tissue section, with single-cell methods resolving the expression profiles and positions of individual cells^5,6^. Manual outlines and landmarks can define anatomical coordinates for cells within individual structures. For example, a villus outline and a landmark at its base define relative cell positions along the crypt-villus axis. However, extending these annotations across many structures or tissue orientations requires repeated manual work. To overcome this limitation, we rationalized that the gene-expression profiles of individual cells and their nearest spatial neighbors could provide a basis for extending annotations of selected structures to all cells throughout the tissue.

We further wondered if these anatomical maps could also support annotation in microscopy, which relies on tissue morphology and the available imaging channels. Spatial transcriptomic datasets pair DAPI images acquired with molecular measurements from the same cells. We reasoned that anatomical maps inferred from those measurements could therefore provide training labels for predicting anatomy from DAPI alone. In principle, this approach could extend to any microscopy data with an acquired or computationally generated DAPI channel, including histology images computationally morphed into DAPI-like images.

To address these tasks, we developed SpatialTRACE (Tissue Region and Axis Coordinate Estimation), comprising SpatialTRACE-Graph and SpatialTRACE-Image. SpatialTRACE-Graph learns from annotations of a few selected structures to predict anatomical axis coordinates or region membership throughout spatial transcriptomic datasets. SpatialTRACE-Image learns from the coordinate and region predictions generated by SpatialTRACE-Graph to map the same anatomy across entire tissue images from DAPI alone. In mouse small intestine, both models predicted crypt-villus and epithelial-distance axis coordinates that closely matched reference maps and distinguished Peyer’s patches from surrounding tissue in sections withheld from supervised training. To test the application of these models outside of spatial transcriptomic datasets, we then fine-tuned SpatialTRACE-Image on annotated immunofluorescence (IF) images to quantify antigen-specific P14 CD8 T cell distributions along these axes and within anatomical compartments. These maps allowed us to compare P14 CD8 T cell abundance at defined anatomical locations between vehicle- and retinoic acid receptor inhibitor-treated sections.

### SpatialTRACE builds anatomical maps from few manually annotated structures

SpatialTRACE starts with selected structures that provide examples of the anatomy to be learned by the model (**Fig. 1a**). For the crypt-villus axis, manual villus outlines and landmarks at the villus base define relative coordinates for cells within each annotated villus. For Peyer’s patch classification, outlines distinguish cells inside the patch from those outside it. These annotations supply training labels without requiring every structure to be annotated (**Fig. 1b**).

**Figure 1.**
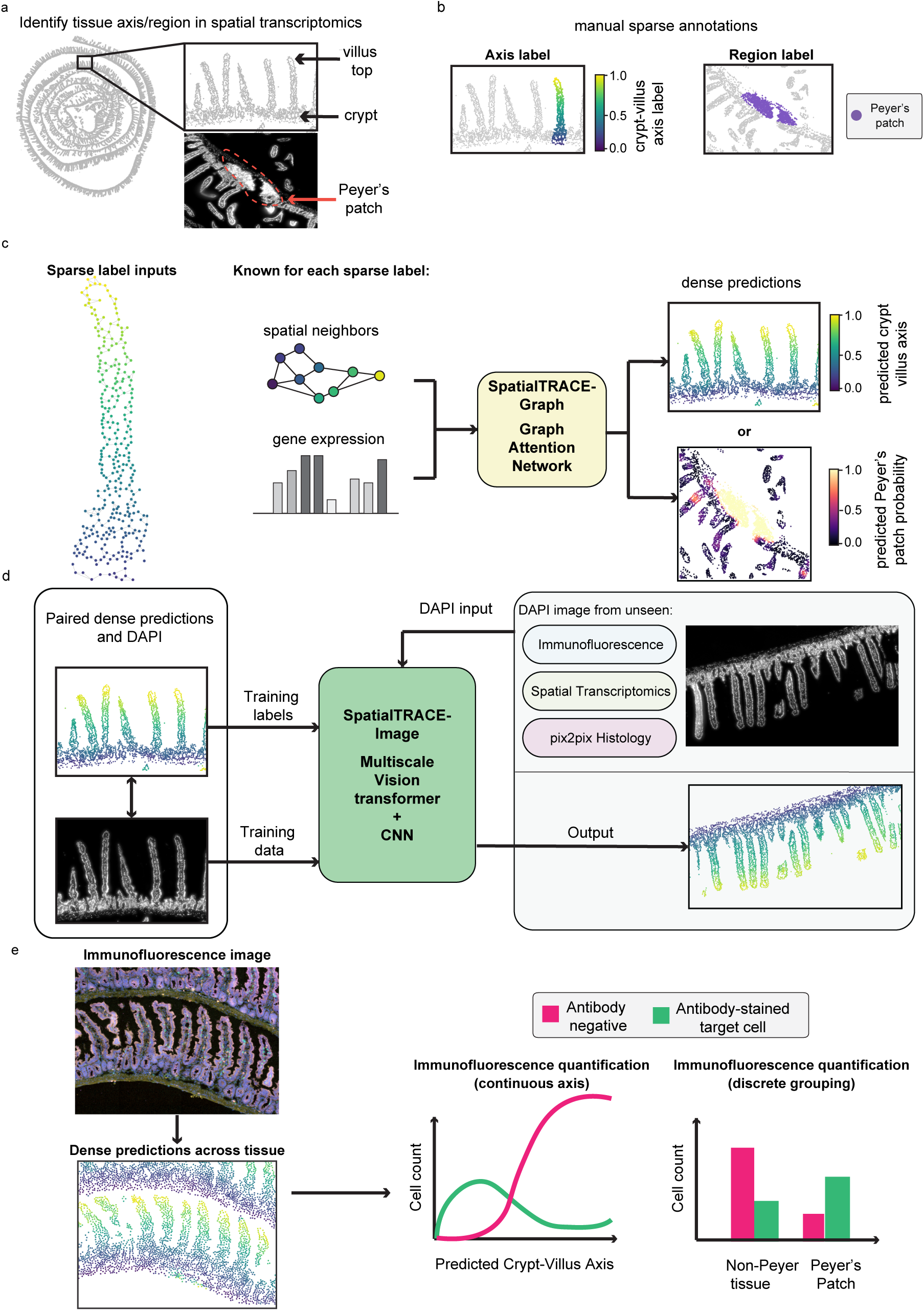
SpatialTRACE creates anatomical maps from manual annotations. **a,** Small-intestine landmarks define the continuous crypt-villus axis and discrete Peyer’s patch regions. **b,** Sparse annotations define coordinates within selected villi and labels for Peyer’s patch classification. **c,** SpatialTRACE-Graph combines gene-expression features with spatial neighborhoods. Sparse labels supervise tissue-wide coordinate or region prediction. **d,** SpatialTRACE-Image learns graph-derived coordinates from matched DAPI images. From left to right: graph-derived reference, matched DAPI image, a separate DAPI field, and its image-model prediction. Axis or region predictions can be made across any imaging data with a DAPI channel. **e,** Schematic illustrating possible applications of SpatialTRACE to immunofluorescence (IF) data, including quantification of antibody-defined cell populations along continuous anatomical axes or within discrete tissue regions.

SpatialTRACE-Graph learns the relationship between these labels and the gene-expression profiles of individual cells and their nearest spatial neighbors. It then uses these relationships to predict anatomical axis coordinates or region membership for all cells across annotated and unannotated tissue sections (**Fig. 1c**). The resulting maps support analysis of gene expression and cellular composition along interpretable anatomical axes and within defined tissue regions.

To extend these anatomical maps beyond spatial transcriptomics, coordinate and region predictions generated by SpatialTRACE-Graph are paired with matched DAPI images to train SpatialTRACE-Image. This model predicts the same anatomical features across entire tissue sections from DAPI alone, without requiring gene-expression measurements from each imaged section (**Fig. 1d**). We chose DAPI because it is widely used in fluorescence microscopy, allowing SpatialTRACE-Image to extend anatomical coordinate and region prediction beyond spatial transcriptomics to a broad range of DAPI-containing imaging datasets.

These image-based anatomical maps can then be used to examine where cell populations localize and how their distributions differ between conditions (**Fig. 1e**). For example, cell abundance can be quantified along the crypt-villus axis or compared inside and outside Peyer’s patches. With DAPI as a shared anchor, SpatialTRACE connects measurements from different imaging modalities to the same anatomical axes and regions.

### SpatialTRACE-Graph predicts continuous coordinates and discrete intestinal regions

SpatialTRACE-Graph uses gene-expression profiles and spatial neighborhoods as inputs to a graph attention network that learns anatomical coordinates or region membership (**Fig. 2a**). Single-cell variational inference (scVI), fitted jointly across sections, compresses gene-expression profiles into 30-dimensional latent representations^7^. Cells are connected to their 20 nearest spatial neighbors within each section. Two attention layers learn how strongly to weight neighboring-cell features^8^, and a task-specific output produces the coordinate or region prediction.

**Figure 2.**
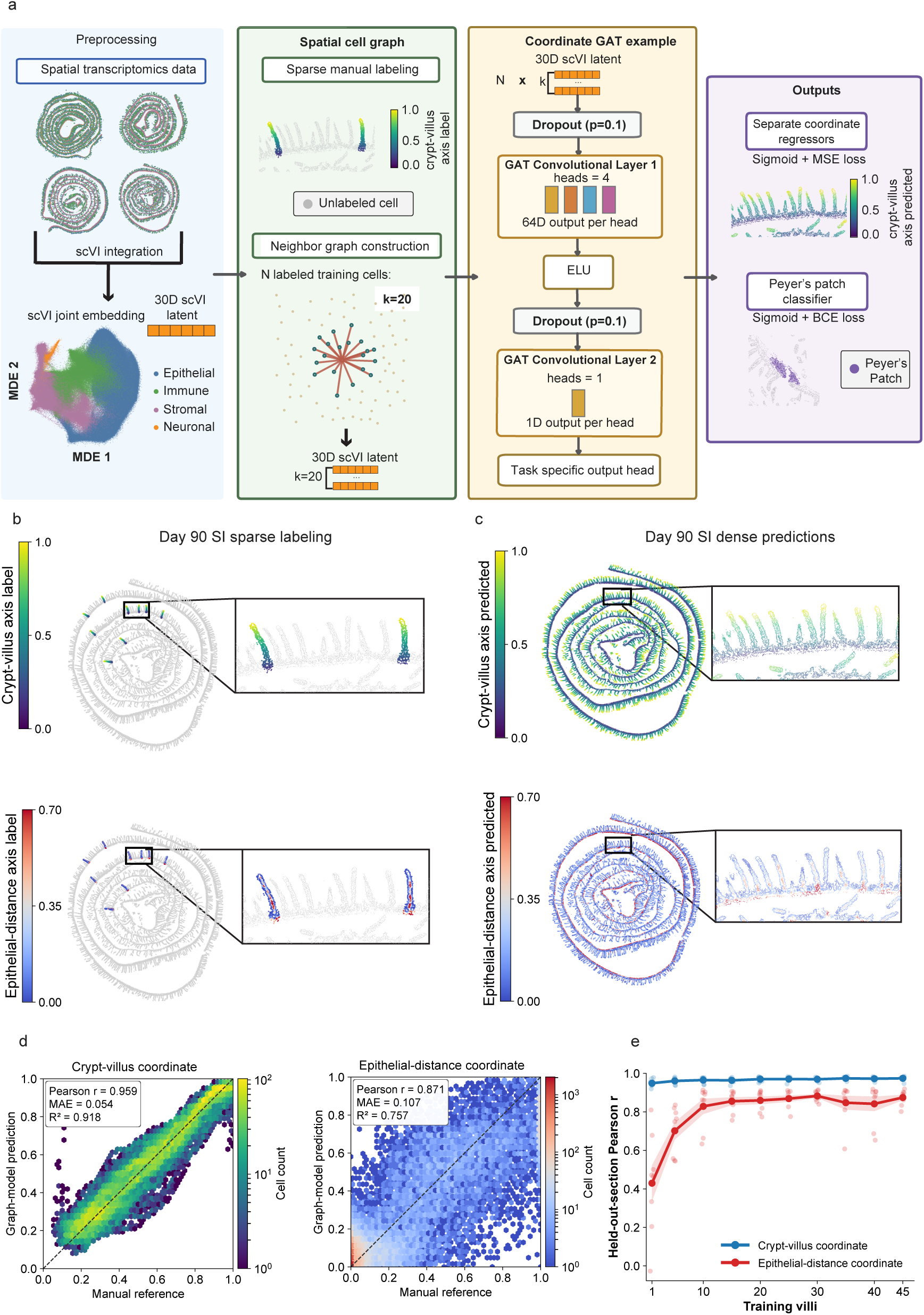
SpatialTRACE-Graph predicts tissue coordinates from sparse annotations. **a,** SpatialTRACE-Graph workflow using gene-expression features learned by single-cell variational inference (scVI) and within-section spatial neighborhoods. Tissue maps and the minimum-distortion embedding (MDE) are colored by cell class. **b,** Sparse references in annotated day 90 villi. Crypt-villus values derive from manual polygons and base landmarks; epithelial-distance values are computed for the same cells. **c,** Dense coordinate maps from SpatialTRACE-Graph models fitted to all sparse labels. **d,** Pooled out-of-fold predictions for 25,436 reference cells from 72 villi across eight sections. Each fold uses six sections for training, one for validation, and one for testing. Pearson correlation (r), mean absolute error (MAE), and the coefficient of determination (R^2^) are shown. Hexagon color shows cell count. **e,** Performance with 1–45 training villi per fold. Points show held-out sections; lines and bands show mean ± standard error of the mean (s.e.m.) across eight sections. Blue and red denote crypt-villus and epithelial-distance prediction, respectively. In **d** and **e**, supervised training, validation, and testing use separate sections. scVI features were learned jointly across the dataset.

To determine whether sparse anatomical annotations could be extended across tissue sections, we applied SpatialTRACE-Graph to mouse small-intestine Xenium data collected 6, 8, 30, and 90 days after LCMV Armstrong infection, an acute systemic viral infection model, with two sections per time point^1^. LCMV Armstrong is a robust and well-characterized immunological model system to study adaptive immune responses in the intestine^9^. Annotations of 72 villi across the eight sections of this dataset provided crypt-villus and epithelial-distance reference coordinates for 25,436 cells (1.2% of the dataset) (**Fig. 2b**). Using these examples as supervision, SpatialTRACE-Graph generated coordinate maps for all cells, reproducing crypt-to-tip gradients and variation in epithelial distance throughout the tissue sections (**Fig. 2c; Extended Data Fig. 1a**).

To test whether these predictions generalized across sections, we held out each of the eight sections in turn for testing, using six other sections for training and one for validation. Across the pooled test cells, predictions were strongly correlated with reference coordinates for both the crypt-villus axis (Pearson *r* = 0.959) and the epithelial-distance axis (*r* = 0.871) (**Fig. 2d**).

We next asked how many annotated training villi were required to recover these coordinates. We varied the number of training villi from 1 to 45 while keeping the validation and test sections fixed. With 10 annotated training villi, our model reached a mean correlation with the reference labels of the held-out-section of approximately 0.97 for crypt-villus position and 0.83 for epithelial distance. Crypt-villus prediction was strong even with one annotated training villus, whereas epithelial-distance prediction improved most over the first 10 villi, with smaller gains from additional annotations (**Fig. 2e; Extended Data Fig. 1c,d**). These findings indicate that SpatialTRACE-Graph can recover anatomical coordinates in held-out tissue sections using as few as 10 annotated training villi.

To assess how spatial neighborhoods and learned attention contributed to this performance, we compared SpatialTRACE-Graph with three alternatives using the same gene-expression features, annotations, and section splits: (1) A cell-only predictor used gene-expression features from individual cells without information from neighbors, (2) a uniform-neighbor model retained spatial connections but weighted neighboring-cell features equally and (3) a randomized-connection model disrupted spatial relationships while preserving the number of neighbors per cell. These comparisons tested whether spatially matched neighbors improved prediction and whether learning their relative contributions provided additional benefit over equal weighting. For crypt-villus position, SpatialTRACE-Graph and the uniform-neighbor model achieved similar mean held-out-section correlations of 0.971 and 0.970, respectively, exceeding the cell-only predictor (0.906) and randomized connections (0.722). Learned attention retained this strong crypt-villus performance while increasing epithelial-distance correlation from 0.677 with equal weighting to 0.871. The cell-only predictor achieved a slightly higher epithelial-distance correlation of 0.900, whereas its crypt-villus correlation remained below that of SpatialTRACE-Graph (**Extended Data Fig. 1e**). Comparisons across matched subsets of 1–45 training villi showed the same overall pattern (**Extended Data Fig. 1f**). Thus, spatial neighborhoods improved crypt-villus prediction relative to cell-only and randomized models, whereas learned attention improved epithelial-distance prediction relative to uniform neighbor weighting.

These differences are consistent with the two axes varying over different physical scales. A nearest-neighbor set may span much of the relatively narrow villus width, including cells in the lamina propria and epithelial cells on opposite sides. Equal weighting could therefore mix features from cells spanning much of the epithelial-distance range, while learned attention can assign different weights to these neighbors. By contrast, the same neighborhood can still provide spatial context along the longer crypt-villus axis. Individual-cell gene expression can also identify epithelial cells, which were used to construct the epithelial-distance reference^1^, potentially contributing to the strong performance of the cell-only predictor.

Finally, to determine whether the same framework could also predict discrete anatomical regions, we trained SpatialTRACE-Graph to classify Peyer’s patch association using manually outlined patches (**Extended Data Fig. 2a,b**). In two sections excluded from supervised training, SpatialTRACE-Graph localized annotated patches and achieved balanced accuracies of 0.91 and 0.97 (**Extended Data Fig. 2c,d**). Thus, sparse annotations supported prediction of both continuous anatomical coordinates and discrete regional identity in additional tissue sections.

### SpatialTRACE-Image predicts anatomical coordinates and regions from DAPI

We next sought to broaden the application of SpatialTRACE beyond spatial transcriptomics to other imaging modalities. Because spatial transcriptomic datasets pair gene-expression measurements with DAPI images from the same cells, we reasoned that coordinates predicted by SpatialTRACE-Graph could provide training labels for an image-based model. We therefore developed SpatialTRACE-Image, which learns from coordinate predictions generated by SpatialTRACE-Graph and the matched DAPI images (**Fig. 3a**). Once trained, SpatialTRACE-Image can predict the same anatomical coordinates from DAPI alone, providing a route to apply these maps to other imaging modalities, including immunofluorescence (IF). To capture the local anatomical context of each cell, SpatialTRACE-Image uses three DAPI crops centered on the same cell at different scales. A fine crop approximately 42 µm wide captures nuclear detail, while local and context crops approximately 166 and 666 µm wide capture neighboring cells and broader tissue organization, respectively. A shared vision transformer processes the local and context crops^10^, with a scale embedding telling the model whether each crop is local or context. Features from the central patches are averaged to represent the area around the target cell. A separate convolutional network processes the fine crop. SpatialTRACE-Image combines features from all three views to predict both anatomical axis coordinates.

**Figure 3.**
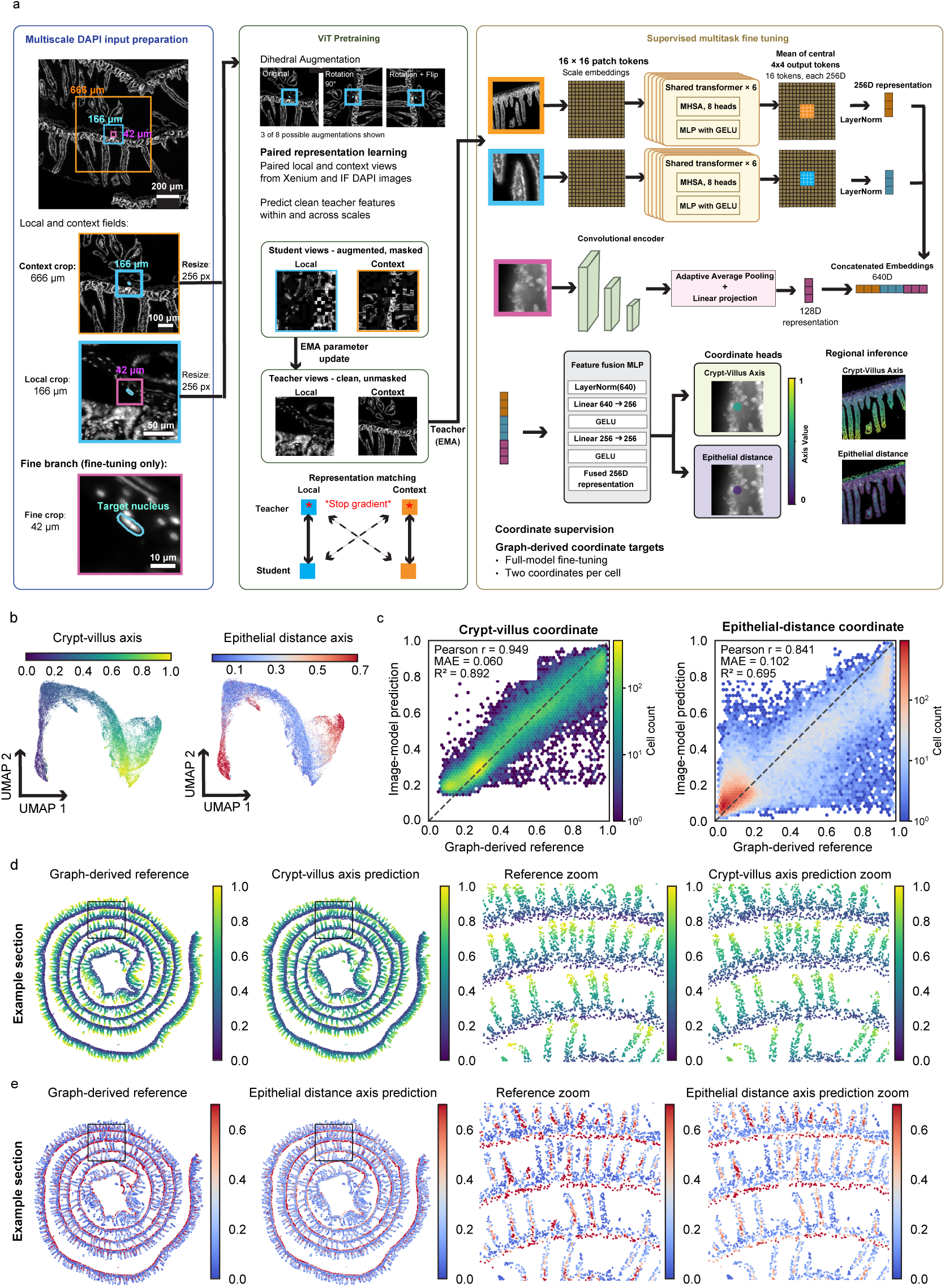
SpatialTRACE-Image learns from graph-based predictions to predict tissue coordinates from DAPI alone. **a,** Model workflow using IF and Xenium DAPI crops for pretraining and Xenium crops for supervised fine-tuning. A vision transformer (ViT) encodes local and context views. The views share encoder weights, with learned scale embeddings identifying each field of view. During pretraining, a student predicts clean teacher features from augmented, masked inputs within and between scales. Teacher weights follow an exponential moving average (EMA) of student weights. Stop-gradient keeps teacher features fixed during each student gradient update. During supervised fine-tuning, central patch features from both scales and convolutional features from a fine-resolution crop predict both coordinates. Crop fields are labeled by physical width; resized network inputs are labeled in pixels. Colored cell-centroid markers show Xenium predictions. MHSA, multihead self-attention; MLP, multilayer perceptron; GELU, Gaussian error linear unit. **b,** Uniform manifold approximation and projection (UMAP) of image embeddings for 30,000 cells, colored by reference coordinates. **c,** Predictions and reference coordinates for 50,000 cells from the same section. Pearson correlation (r), mean absolute error (MAE), and coefficient of determination (R^2^) are shown. Hexagon color shows cell count; dashed lines indicate equal values. **d,e,** Full-section reference, prediction, reference zoom, and prediction zoom for crypt-villus position (**d**) and epithelial distance (**e**). Full maps include all 50,000 cells. References are SpatialTRACE-Graph coordinates. Panels **b–e** show a day 8 replicate section excluded from supervised training and validation.

Before supervised fine-tuning, the vision transformer learns image features from unlabeled Xenium and IF DAPI crops through paired representation pretraining. A student network predicts features from augmented, partly masked local and context crops, while a teacher network provides targets from the corresponding clean images^11–13^. Teacher weights follow an exponential moving average of student weights, while stop-gradient blocks backpropagation through the teacher^12^. Matching student and teacher features within and across the two scales provides supervision without anatomical labels (**Extended Data Fig. 3a,b**). The pretrained transformer then initializes full-model fine-tuning on coordinates predicted by SpatialTRACE-Graph (**Extended Data Fig. 3c**).

In a day 8 Xenium section, image representations varied with both crypt-villus position and epithelial distance (**Fig. 3b**). Across 50,000 evaluated cells, predictions were strongly correlated with the references generated by SpatialTRACE-Graph for crypt-villus position (*r* = 0.949) and epithelial distance (*r* = 0.841) (**Fig. 3c**). The predicted maps reproduced repeated crypt-villus axis structure and variation in epithelial distance across the section (**Fig. 3d,e**). Thus, DAPI images captured sufficient anatomical organization for SpatialTRACE-Image to recover much of the spatial organization captured in the transcriptomic reference maps.

We next asked whether pretraining improved coordinate prediction when fewer labeled image examples were available for supervised fine-tuning. For this comparison, we deliberately restricted fine-tuning to the same 4,084 cells, representing 5% of the coordinate-labeled training set, and compared random initialization with masked-pixel reconstruction and paired representation pretraining. This reduced-supervision comparison was separate from training the main coordinate model, which used the full training set. Across three matched runs, paired representation pretraining modestly increased mean crypt-villus correlation from 0.860 to 0.875 and mean epithelial-distance correlation from 0.692 to 0.718 relative to random initialization. Crypt-villus correlation improved in all three runs, and epithelial-distance correlation improved in two of three. Masked-pixel reconstruction produced intermediate mean correlations (**Extended Data Fig. 3d**). These results indicate that paired representation pretraining modestly improved coordinate prediction when supervised training data were limited.

To determine which image scales contributed to the trained predictions, we shuffled one crop type among cells while keeping the remaining inputs and model weights fixed. This disrupted the correspondence between a view and the cell being predicted without removing the view from the model. Shuffling each of the three crop types reduced correlation for both coordinates, indicating that the trained model used information from all three views. Local-crop shuffling caused the largest reductions, followed by context-crop shuffling. Fine-crop shuffling had smaller effects but still reduced both correlations (**Extended Data Fig. 3e**).

We next removed each crop branch and retrained the remaining architecture to test whether the views contributed complementary information. Removing the fine branch reduced correlations for both coordinates, supporting the use of fine-scale features alongside the larger fields of view. Removing the local branch reduced epithelial-distance correlation while leaving crypt-villus prediction nearly unchanged. The context branch contributed most clearly to crypt-villus prediction: removing it reduced crypt-villus correlation, although epithelial-distance correlation improved. Retaining all three views yielded the highest crypt-villus correlation and strong epithelial-distance performance, combining fine-scale information with local and tissue-level context (**Extended Data Fig. 3f**).

Next, we tested whether SpatialTRACE-Image could distinguish discrete anatomical regions. We trained a Peyer’s patch classifier using the same pretrained encoder and three-branch architecture, with manual patch labels or SpatialTRACE-Graph probabilities used as supervision. Predictions matched the manually annotated patch in an example Xenium section excluded from supervised training (**Extended Data Fig. 3g**). Across 819 cells from two sections excluded from supervised training, fine-tuning all model components improved AUROC from 0.87–0.88 with a fixed transformer to 0.93–0.94, indicating strong discrimination between patch and surrounding tissue (**Extended Data Fig. 3h**). Thus, SpatialTRACE-Image could recover both continuous anatomical coordinates and discrete regional identity from DAPI images, allowing cellular measurements to be placed within both anatomical axes and defined tissue regions.

### SpatialTRACE-Image maps intestinal T cell distributions after retinoic acid receptor inhibition

We next asked whether SpatialTRACE-Image could generalize from Xenium-associated DAPI images to previously unseen IF images to recover intestinal anatomical axes. Retinoic acid promotes intestinal T cell homing and supports the formation and maintenance of intestinal tissue-resident memory T cells^14–16^. To track an antigen-specific CD8 T cell response, we used the P14 T cell receptor transgenic model, in which congenically marked P14 CD8 T cells recognize the LCMV GP33 epitope and can be followed after adoptive transfer and infection. After infection, mice were either treated with a retinoic acid receptor inhibitor (RARi) or dimethyl sulfoxide (DMSO) as a control. Small-intestine IF images were collected on day 7 after acute systemic LCMV Armstrong infection. CD45.1 and CD8α staining identified P14 CD8 T cells, and E-cadherin marked the epithelium (**Fig. 4a**). We then applied SpatialTRACE-Image to examine the anatomical distribution of P14 CD8 T cells in these sections.

**Figure 4.**
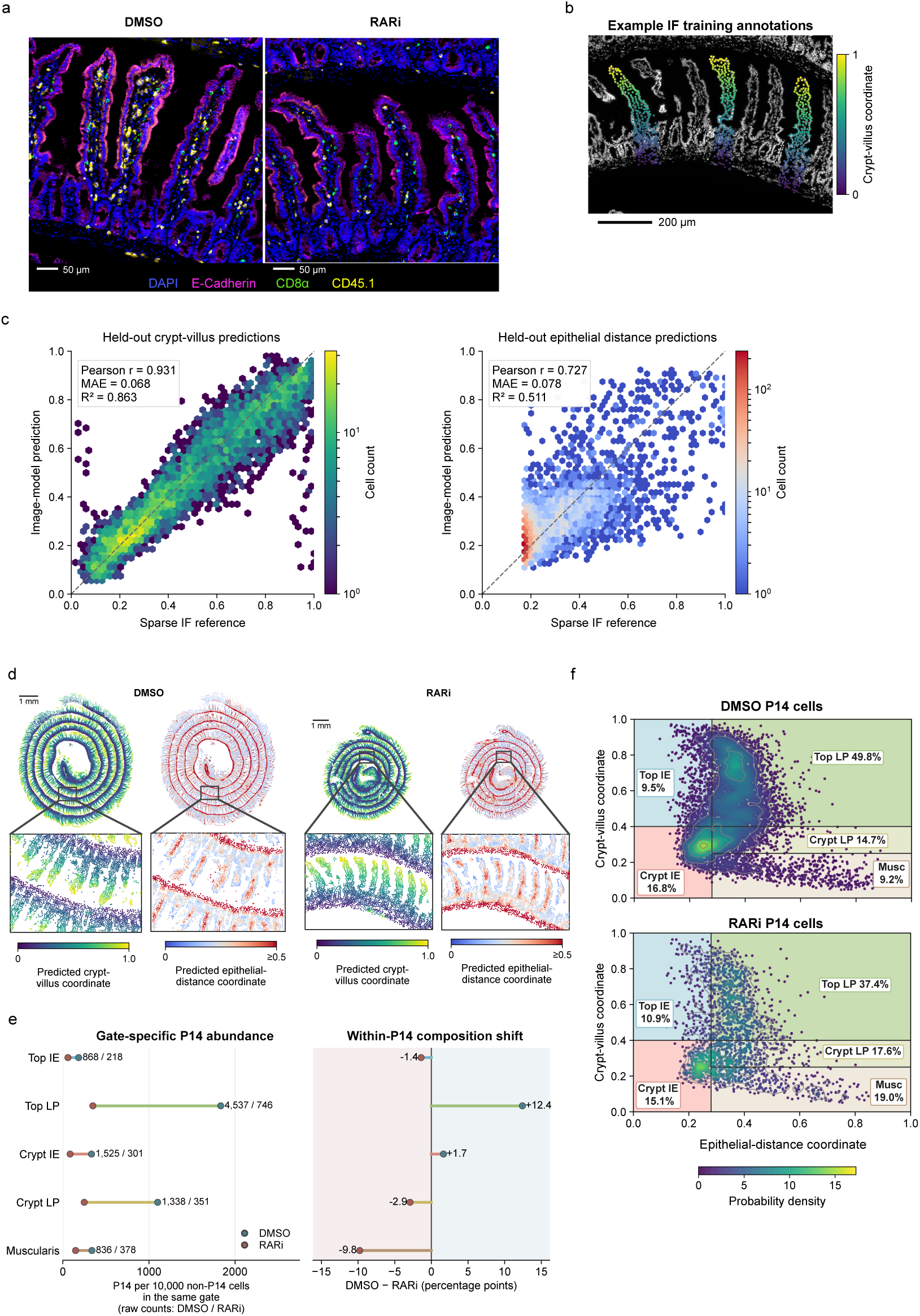
SpatialTRACE-Image maps P14 cell abundance and anatomical distribution in IF. **a,** Multiplex IF images of small intestine on day 7 of lymphocytic choriomeningitis virus (LCMV) Armstrong infection, comparing dimethyl sulfoxide (DMSO) vehicle and retinoic acid receptor inhibitor (RARi) treatment. Colors show DAPI (blue), E-cadherin (magenta), CD8α (green), and CD45.1 (yellow). CD8α and CD45.1 coexpression identifies transferred P14 CD8^+^ T cells. Scale bars, 50 µm. **b,** Example crypt-villus references for 749 training cells in three villi, overlaid on DAPI. Scale bar, 200 µm. **c,** IF-model predictions for 4,843 cells from 19 test villi, with all three crop scales spatially disjoint from training and validation. Crypt-villus references derive from manual villus annotations; epithelial-distance references derive from QuPath epithelial classifications and a common training-derived scale. Pearson correlation (r), mean absolute error (MAE), and coefficient of determination (R^2^) are shown. Hexagon color shows cell count; dashed lines indicate equal values. **d,** Whole-section predictions and magnified boxed regions. Whole-section scale bars, 1 mm. **e,** P14 cells per 10,000 non-P14 cells in the same gate (left) and within-P14 composition differences (right; DMSO minus RARi percentage points). Raw P14 counts are shown beside each gate. **f,** Immune Allocation Maps (IMAPs) of 9,104 DMSO and 1,994 RARi P14 cells. Gate labels show percentages of P14 cells. Gate names describe anatomical interpretations of the coordinate ranges. Musc, muscularis; IE, intraepithelial; LP, lamina propria. Panels **d–f** compare one section per condition descriptively.

We fine-tuned the Xenium-trained SpatialTRACE-Image model using annotated IF villi (**Fig. 4b; Extended Data Fig. 4a**). Before IF-specific fine-tuning, the Xenium-trained model already predicted crypt-villus and epithelial-distance coordinates from IF images, with Spearman correlations of 0.74 and 0.40, respectively. Fine-tuning increased these correlations to 0.94 and 0.67, showing that anatomical prediction generalized to IF in a zero-shot setting and improved further with IF-specific supervision (**Extended Data Fig. 4b,c**).

Evaluation used 4,843 cells from 19 villi, with crops at all three scales spatially disjoint from training and validation. Crypt-villus references were derived from manual villus annotations, while epithelial-distance references were calculated from E-cadherin-based epithelial classifications as described in **Methods**. Fine-tuned predictions closely followed the manual crypt-villus coordinates (Pearson *r* = 0.931) and captured epithelial-distance variation (*r* = 0.727) (**Fig. 4c**). We used the IF-fine-tuned model for all subsequent coordinate maps and P14 analyses.

SpatialTRACE-Image then assigned crypt-villus and epithelial-distance coordinates to cells throughout each IF section (**Fig. 4d**). To convert these continuous coordinates into anatomically interpretable compartments, we applied fixed coordinate gates that defined upper-villus and crypt-associated intraepithelial regions (Top IE and Crypt IE), the corresponding lamina propria regions (Top LP and Crypt LP), and the muscularis. We projected the resulting gate assignments back onto the IF images to relate these coordinate-defined compartments to visible tissue structure (**Extended Data Fig. 4d**).

For each compartment, we quantified P14 CD8 T cell abundance relative to non-P14 cells in the same compartment and the fraction of the total P14 population located there. The RARi section contained fewer detected P14 CD8 T cells overall, with lower normalized abundance in all five compartments (**Fig. 4e**). To visualize how P14 CD8 T cells were distributed across intestinal anatomy, we used Immune Allocation Maps (IMAPs), which position cells in a two-dimensional coordinate space defined by the predicted crypt-villus and epithelial-distance axes^1^. Using the same anatomical boundaries between conditions, IMAPs revealed a redistribution of the P14 population after RAR inhibition. The fraction of P14 cells in the upper-villus lamina propria decreased from 49.8% in DMSO to 37.4% in RARi, whereas the fraction in the muscularis increased from 9.2% to 19.0%. P14 cell frequencies in the other three compartments remained similar. Importantly, the enrichment in the muscularis reflected a larger relative share of the P14 population rather than an absolute increase in local P14 cell abundance (**Fig. 4f**).

To assess whether these differences reflected broader differences in tissue composition, we mapped non-P14 cells using the same axes and gates. Non-P14 gate fractions were similar between sections, in contrast to the larger P14 differences in upper-villus lamina propria and muscularis. These results indicate that the observed redistribution was concentrated in the P14 population rather than reflecting a comparable shift in the tissue-wide reference population (**Extended Data Fig. 4e**).

Lastly, we also applied the Xenium-trained region classifier directly to IF DAPI images without additional supervised IF training. In both sections, high predicted Peyer’s patch probabilities coincided with compact regions that visually resembled Peyer’s patches, showing agreement between zero-shot regional predictions and tissue morphology (**Extended Data Fig. 4f**).

Together, these analyses show that SpatialTRACE-Image can predict both continuous anatomical coordinates and discrete regions in IF data. SpatialTRACE connects cell measurements to tissue anatomy and resolves compartment-specific distribution differences that whole-section counts alone would miss.

## Discussion

We developed SpatialTRACE as a general framework for extending sparse anatomical annotations into tissue-wide maps that can be shared across spatial transcriptomics and microscopy. SpatialTRACE-Graph establishes these maps within spatial transcriptomics, and SpatialTRACE-Image extends anatomical prediction to microscopy data. In IF, these coordinates resolved compartment-specific differences in P14 CD8 T cell distributions that whole-section counts did not capture.

SpatialTRACE-Graph uses learned attention to retain useful spatial context without weighting all neighboring cells equally. The component comparisons indicate that spatial neighborhoods are particularly informative for crypt-villus position, whereas learned attention improves epithelial-distance prediction relative to uniform neighbor weighting. For SpatialTRACE-Image, shuffling and branch ablations indicate that the three views contain complementary but partly overlapping anatomical information. Retraining allows the remaining views to partially compensate for some of the information lost when a branch is removed. Together, these analyses suggest that anatomical position is encoded across multiple spatial scales rather than within a single local image feature.

SpatialTRACE-Image recovers anatomical axes and regions that have corresponding patterns in DAPI, whether in nuclear morphology or tissue organization. The zero-shot IF predictions show that these patterns generalize across imaging datasets. However, additional fine-tuning strongly improves coordinate recovery. Our data show that DAPI can provide a common anatomical reference that connects spatial transcriptomic measurements with downstream microscopy datasets.

Applied to the RARi IF section, SpatialTRACE-Image revealed relative depletion of P14 CD8 T cells from upper-villus lamina propria and enrichment in the muscularis. Non-P14 cells did not show comparable shifts, arguing against sampled tissue composition as the sole explanation. This pattern suggests that retinoic acid signaling may influence intratissue positioning in addition to overall intestinal accumulation. Given the established roles of retinoic acid in intestinal homing and tissue-resident memory formation^14–16^, altered access to or persistence within specific compartments could connect spatial environments encountered during early localization to later residency.

Pairing spatial transcriptomics with SpatialTRACE-Graph provides an efficient way to generate large sets of anatomical training labels from a few annotated structures. Matched DAPI images turn these labels into training data for SpatialTRACE-Image, which can then predict the same anatomy across large microscopy datasets. DAPI provides an anchor for fluorescence microscopy, and image translation could extend that anchor to conventional histology and archived tissue collections when nuclear organization is preserved^17,18^. More broadly, consistent anatomical definitions could allow axes defined in spatial transcriptomics to serve as a shared coordinate system across imaging modalities.

## Methods

### Mice, infection, adoptive transfer, and inhibitor treatment

Mice were on a C57BL/6J background and were bred at the University of California, San Diego (UCSD), or purchased from The Jackson Laboratory. Animal studies were approved by the UCSD Institutional Animal Care and Use Committee and followed UCSD guidelines.

CD45.1 P14 donor cells (50,000 per mouse) were transferred intravenously by retro-orbital injection into CD45.2 recipient mice one day before infection. Recipient mice were infected intraperitoneally with 2 × 10^5^ plaque-forming units of lymphocytic choriomeningitis virus, Armstrong strain (LCMV Armstrong).

For immunofluorescence (IF) analysis, mice received 20 µg retinoic acid receptor-α inhibitor (RARi; ER 50891, Tocris) dissolved in DMSO, or DMSO control, from days 0 to 7 of infection. Small intestines were collected on day 7. One section per condition was analyzed (**Figure 4**).

### Tissue preparation, immunofluorescence staining, and imaging

Small-intestine tissues were fixed overnight at 4 °C with 4% paraformaldehyde, embedded in optimal cutting temperature compound, and cryosectioned at 10 µm. Sections were blocked with Dako blocking reagent for 1 h. Antibodies were diluted in Dako antibody diluent and incubated overnight at 4 °C. Sections were stained for CD45.1, E-cadherin, and CD8α, counterstained with DAPI, and mounted with Vectashield Vibrance. Multichannel IF images were acquired at 40× on an Olympus VS200 Slide Scanner (UCSD Microscopy Core) or a Zeiss LSM700 confocal microscope.

Cell segmentation and classification were performed in QuPath^19^. DAPI-based cell detection used a requested pixel size of 0.5 µm, background radius of 8 µm, Gaussian sigma of 1.5 µm, nuclear area range of 10–400 µm^2^, and intensity threshold of 100. Detection included opening by reconstruction, shape-based splitting, and boundary smoothing. Cell boundaries extended 5 µm beyond the detected nuclei.

QuPath classified epithelial and non-epithelial cells using E-cadherin signal. A trained classifier identified transferred P14 CD8^+^ T cells as CD45.1 and CD8α double-positive cells.

### Spatial transcriptomics data

Xenium spatial transcriptomics data came from a published study that described tissue preparation, panel design, imaging, and segmentation^1^. Analyses used AnnData objects containing the published cell-by-gene matrices and spatial coordinates, filtered as reported in that study^1,20^.

Graph inputs comprised a 30-dimensional scVI representation, cell spatial coordinates, and section identifiers^7^. A two-dimensional minimum-distortion embedding and cell-class annotations were used for visualization (**Figure 2a**)^21^. The scVI representation was learned jointly across sections and then fixed.

### Sparse anatomical reference coordinates

Crypt-villus references were derived from villus polygons and crypt-side base keypoints annotated in LabelMe^22^. Each cell within a polygon was assigned its Euclidean distance from the base, divided by the maximum distance among cells in that villus. Values ranged from 0 near the base to 1 at the most distant cell. Image annotations were transformed into cell-coordinate space.

The Xenium annotation set contained 25,436 labeled cells in 72 villi across eight sections sampled on days 6, 8, 30, and 90, with two sections per time point. The section labels replicate 1 and replicate 2 distinguish these two sections at each time point. IF villus polygons and base keypoints were annotated separately and matched to cells by their coordinates and object identifiers.

The upstream Xenium pipeline calculated a dimensionless epithelial-proximity reference within each source section. It divided the mean distance to the five nearest cells annotated as epithelial by the mean distance to the five nearest cells overall; the query cell was included when present in a reference tree. The ratio was divided by its section-wide 99th percentile and clipped to [0, 0.6]. Within each manually annotated villus, these values were then min–max rescaled to [0, 1].

IF epithelial-distance references used QuPath epithelial classifications. Within each tissue cluster, the mean distance to the five nearest epithelial cells was divided by the mean distance to the five nearest cells overall, including the query cell when present. Ratios were scaled by their 99th percentile among the 5,822 training cells. This training-derived scale was applied to both sections and all partitions, followed by clipping to [0, 1]. Xenium and IF coordinates were calibrated separately.

### Graph attention networks for coordinate prediction

Separate graphs connected cells within each section to their 20 nearest spatial neighbors, with reciprocal edges added and duplicates removed. Each node carried its 30-dimensional scVI vector. Each training batch contained 1,024 labeled cells, termed seed cells, and 10 sampled neighbors at each of two graph hops. Loss was calculated on the seed cells.

Separate models predicted crypt-villus and epithelial-distance coordinates. Each contained two graph attention layers^8^. The first produced 64 features per head using four heads, followed by exponential linear unit (ELU) activation^23^ and dropout of 0.1. The second mapped the concatenated 256-dimensional representation to one scalar using a single head. A sigmoid constrained predictions to [0, 1].

Graph-coordinate training minimized mean squared error using the Adam optimizer^24^ with learning rate 10^-3^ and weight decay 10^-5^. Training lasted up to 500 epochs, with early-stopping patience of 100 epochs.

Graph-coordinate prediction was evaluated in eight section-disjoint folds, with six sections for training, one for validation-based checkpoint selection, and one for testing (**Figure 2d**). Each section served as the test once, yielding one out-of-fold prediction per labeled cell. Inference used all neighbors at both hops. Metrics were calculated for the pooled 25,436 cells and separately by held-out section.

Annotation requirements were evaluated using the same section split with nested training subsets of 1 villus and multiples of 5 through 45 villi, selected in a fixed random order from the six training sections (**Figure 2e**). Validation and test sections remained unchanged across training-set sizes. Curves show the mean and standard error across eight held-out sections.

The fitting-time and additional error analyses used the same eight section-disjoint folds and nested training-villus subsets (**Figure 2e; Extended Data Figure 1b–d**). MAE and R^2^ were summarized by the mean and standard error across held-out sections. Fitting time was measured for eight folds, ten training-villus counts, and two coordinates, totaling 160 fits on an NVIDIA RTX A6000. Timing covered model fitting, validation, and checkpoint writing. Median and interquartile range summarized fitting time across folds. Villus counts refer to training annotations; each fold also used one annotated section for validation.

Dense coordinate maps were generated using models fitted to pooled labels from eight sections, with 90% of labeled cells for training and 10% for validation (**Figure 2c; Extended Data Figure 1a**). The resulting predictions provided tissue-wide coordinate maps and training targets for the image model.

### Graph component and annotation-budget comparisons

The full graph model was compared with a cell-only predictor, a uniform-neighbor graph, and a graph with randomized connections (**Extended Data Figure 1e**). All models used the same 30-dimensional scVI features. The cell-only predictor used two linear layers with 256 hidden units, ELU activation, dropout, and a sigmoid output. Uniform-neighbor models replaced learned attention with uniform aggregation while retaining the graph architecture, linear transformations, self-loops, and dropout.

Randomized graphs permuted cell identities within each section among cells with the same number of neighbors. Applying the permutation to both endpoints of every edge preserved the number of neighbors for every cell and reciprocal connections while disrupting their relationship to cell location. Connections were fixed before training and retained for validation and testing. Each comparator was fitted separately for each coordinate and fold using the same annotations, section partitions, optimization settings, and validation-loss selection as the full model.

The full graph, cell-only, and uniform-neighbor models were compared using the nested training-villus subsets (**Figure 2e; Extended Data Figure 1f**). All three models used identical annotated villi at each count, with validation and test sections unchanged. Full-graph results were compared with separately trained cell-only and uniform-neighbor models at every count. Pearson correlation and MAE were calculated by held-out section. Lines and bands show the section mean and standard error across eight sections.

### Peyer’s patch graph classifier

A binary graph attention network classified Peyer’s patch membership using 30-dimensional scVI features and 20-neighbor graphs constructed separately within each section. Cells within manually annotated Peyer’s patch polygons were labeled positive, and cells outside the polygons were labeled negative. Training sampled 25 negative cells per positive cell. The annotation set contained 7,882 positive cells among 2,075,222 cells in eight sections.

Training used replicate-1 sections from days 6, 8, 30, and 90 of infection. Validation used replicate-2 sections from days 6 and 8, and testing used replicate-2 sections from days 30 and 90. Training included 2,509 positive and 62,725 negative seed cells; validation and testing included 1,896 and 3,477 positive cells, respectively. The two test sections contained 500,050 cells. scVI features and feature normalization were learned jointly across the dataset. Predictions from these test sections were inspected against manual patch outlines during classifier development.

The first attention layer produced 128 features per head using four heads; the second produced 128 features using one head, followed by a linear binary classifier. ELU activations and dropout of 0.15 were used between layers. Training minimized binary cross-entropy with logits and a positive-class weight of 25. AdamW^25^ used learning rate 10^-4^, weight decay 10^-4^, gradient-norm clipping at 1, and batches of 4,096 seed cells. Neighbor sampling included 15 and 10 cells at the first and second hops, respectively. Training allowed up to 80 epochs with early-stopping patience of 12 epochs. Validation average precision selected epoch 10 after 22 epochs. A probability threshold of 0.95 maximized validation F1 over a grid from 0.05 to 0.95 in steps of 0.005. This threshold was fixed for testing.

### DAPI inputs and physical crop scaling

Image models used single-channel DAPI crops centered on cell centroids. Local, context, and fine fields were approximately 166, 666, and 42 µm wide, respectively.

Source pixel sizes were approximately 0.213 µm for Xenium and 0.162 µm for the IF images (**Figure 4**). Native local, context, and fine crop widths were 783, 3,132, and 196 pixels for Xenium, and 1,024, 4,097, and 256 pixels for IF. Local and context crops were resized to 256 × 256 pixels, and fine crops to 128 × 128 pixels.

Xenium crop origins were rounded to the nearest pixel at the selected image-pyramid level; IF crop origins were rounded down to the nearest source pixel. Crops were zero-padded at image boundaries. Intensities were normalized using the 1st and 99.8th percentiles, clipped to [0, 1], and resized by bilinear interpolation. Percentiles for large crops used subsampled pixels. For Xenium supervised training and inference, crops were sampled from image-pyramid levels appropriate for the final input resolution, quantized to 8-bit values, and scaled to [0, 1]. IF supervised inputs retained floating-point intensities after normalization and resizing. Pretraining inputs used an intermediate 512 × 512 image before resizing to 256 × 256 pixels, with 8-bit quantization after each step.

### Shared, scale-aware vision transformer architecture

One shared vision transformer ViT encoded local and context crops. Each crop was divided into 256 patches of 16 × 16 pixels, projected to 256-dimensional tokens. A learned CLS token was prepended. Positional embeddings and one of two scale embeddings were added to all tokens. Scale embeddings identified each input as local or context.

Each of the six transformer blocks contained eight attention heads and a feed-forward network with 1,024 hidden units and GELU activation^26^. Layer normalization preceded the attention and feed-forward operations; dropout probability was 0. The shared encoder processed each scale separately. Its output was the average of the central 4 × 4 patch tokens after the final block, followed by layer normalization. The CLS token participated in attention.

The fine CNN used three 3 × 3 convolutions with 32, 64, and 128 channels, GELU activations, and stride 2 in the last two layers. Adaptive average pooling and a linear projection produced 128 features.

Concatenating local, context, and fine features yielded a 640-dimensional vector. Fusion used layer normalization and two 256-unit linear layers, each followed by GELU activation. Separate linear heads with sigmoid outputs predicted each coordinate. The full model contained approximately 5.2 million parameters.

### Paired representation pretraining

Pretraining used unlabeled pairs of local and context crops centered on the same cell: 22,048 pairs from seven Xenium sections and 9,963 from four IF sections. Holding out one section per modality gave 27,458 training and 4,553 validation pairs.

Both student crops received the same geometric and intensity augmentations. Eight combinations of rotations by multiples of 90° and reflections cycled across epochs. Intensity augmentation occurred with probability 0.8 and used contrast 0.7–1.3, brightness within ±0.1, and Gaussian blur sigma 0–1.5 pixels. Patch tokens were independently replaced with a learned mask token^11^ at probability 0.4 at each scale. The teacher received the original paired crops. Validation used the original images with deterministic student masking.

Student and teacher each used the scale-aware ViT with the central patch readout described above. Their projection heads mapped encoder features through layer normalization, a linear layer from 256 to 512 dimensions, GELU activation, and a linear layer back to 256 dimensions. The student additionally used a predictor with the same architecture^12^.

The teacher encoder and projection head were initialized from the student. After each optimization step, teacher weights were updated by an exponential moving average (EMA) of student weights, producing a slowly changing target network. EMA momentum increased from 0.996 toward 1 on a cosine schedule. Stop-gradient blocked backpropagation through the teacher, so teacher parameters changed only through the EMA update.

The pretraining objective matched student predictions to teacher features from the same and opposite scales. Alignment loss was 2 − 2p·z for unit-normalized student and teacher vectors p and z, averaged over both scale assignments. Variance regularization penalized feature standard deviations below 1 to promote variation across cells. Covariance regularization penalized squared off-diagonal covariances to reduce redundant features^27^. Both penalties were averaged across scales. Total loss was the mean of same-scale and cross-scale alignment losses plus five times the variance penalty and 0.04 times the covariance penalty.

Pretraining used the AdamW optimizer for 35 epochs with batch size 16, weight decay 0.04, and cosine learning-rate decay from 2 × 10^-5^ to 2 × 10^-7^ starting at the first epoch. Gradient values and norm were clipped at 1. Minimum validation loss selected epoch 33. The selected teacher encoder, including scale embeddings, positional embeddings, and normalization, initialized supervised training.

### Xenium supervised fine-tuning and evaluation

Supervised training fitted the full three-branch model to coordinate targets generated by SpatialTRACE-Graph. The shared ViT inherited teacher weights; the fine CNN, fusion layers, and prediction heads were initialized anew. Training used images from 81,656 cells in six sections, and validation used 5,099 cells in one section. The image-coordinate development set comprised 5,107 cells from four sections excluded from supervised training and validation.

Training minimized equally weighted Smooth L1 losses for the two coordinates. Epithelial-distance loss was computed for cells with available references. AdamW used learning rate 10^-4^, weight decay 10^-4^, batch size 16, and 20 epochs, with unaugmented inputs. Minimum validation loss across three training runs selected the epoch-9 checkpoint.

Image-model predictions were compared with coordinates generated by SpatialTRACE-Graph for 50,000 cells from the day 8 Xenium section excluded from supervised fitting and validation (**Figure 3b–e**). Predictions and references were matched by cell identity and spatial coordinates. This larger spatial sample illustrated tissue-wide coordinate patterns.

The fused 256-dimensional image representations were visualized by UMAP for a fixed subset of 30,000 cells. UMAP used 30 neighbors, minimum distance 0.15, and a cosine metric. All colorings used the same projection.

Example inference used a day 90 Xenium field (**Figures 1d and 3a**). Paired local and context crops from Xenium and IF DAPI images were used to illustrate pretraining (**Figure 3a**).

### Pretraining and image-branch comparisons

Random initialization, pretraining by masked-pixel reconstruction, and paired representation pretraining were compared under reduced coordinate supervision (**Extended Data Figure 3d**). All three strategies used the same 4,084 coordinate-labeled training cells, approximately 5% of the 81,656 cells in the full coordinate-training set, with unchanged validation and image-coordinate development sets. Each strategy used three matched fine-tuning runs. Architecture, fine-tuning settings, and validation-based checkpoint selection were matched; the pretraining procedures differed. The main coordinate model and branch-retraining comparisons used the full coordinate-training set.

Sensitivity to disrupted input pairing was measured by shuffling inputs at one scale at a time (**Extended Data Figure 3e**). Images at one scale were shuffled among cells in the image-coordinate development set within batches of 128 so that every image was assigned to a different cell. Other inputs, targets, and model weights remained fixed. Five repeats per scale measured the drop in Pearson correlation.

Branch ablations compared the full local, context, and fine model with architectures lacking one crop branch (**Extended Data Figure 3f**). Each reduced architecture was retrained after removal of the local, context, or fine branch. Retained branches used identical starting weights: the transformer inherited the same pretrained encoder, and the fine CNN used the same random initialization. Models used the full coordinate-training set, with matched preprocessing, training settings, and validation-loss checkpoint selection. All four models were evaluated on the same image-coordinate development set using Pearson correlation and MAE.

The branch comparison used the random initialization associated with the full model selected by validation among three runs. Each reduced architecture was fitted once using that matched initialization, giving one training run per architecture.

### Peyer’s patch DAPI classifier

All Peyer’s patch image classifiers used the same shared, scale-aware ViT, central 4 × 4 patch-token readout, fine CNN, and fusion architecture as coordinate prediction. A single sigmoid output predicted region membership. The shared transformer was initialized from the same representation-pretrained EMA teacher, retaining scale and positional embeddings and the CLS token. The fine CNN, fusion layers, and binary output were initialized anew.

Four variants crossed manual binary labels or graph-derived probabilities (soft targets) with a frozen transformer or full-model fine-tuning. The fine CNN, fusion layers, and binary output were trained in both adaptation schedules. All four models shared initial weights, section partitions, and sampled cells. Section assignments matched those of the Peyer’s patch graph classifier.

Training used 70 positive and 210 negative cells, and validation used 55 positive and 431 negative cells. Evaluation used an 819-cell classifier development cohort comprising 91 positive and 728 negative cells from two sections excluded from supervised training (**Extended Data Figure 3h**). Positive cells connected within 80 µm formed regions containing at least 25 positive cells. Sampling selected at most 80 positive cells per region, with at least 40 µm between selected cells. Negative cells were sampled within each section using the same minimum spacing.

Classifier training minimized binary cross-entropy with logits, with positive-class weights of 3 for manual targets and 2.84 for soft targets. AdamW used learning rate 10^-4^, weight decay 10^-4^, batch size 16, and 20 epochs, with unaugmented inputs. Validation average precision selected each checkpoint; all four checkpoints were selected before evaluation. Full-model training on manual labels was specified for regional inference before fitting, and validation selected epoch 20. Binary predictions used a fixed probability threshold of 0.5.

The manual-label, full-model classifier generated predictions for a window from the day 90 Xenium test section and for DMSO and RARi IF windows (**Extended Data Figures 3g and 4f**). The IF windows contained 2,518 DMSO and 2,263 RARi cells and used a shared linear 0–1 probability scale. IF patch status was not independently annotated; these maps therefore provide a qualitative display of classifier output.

### IF fine-tuning and crop-disjoint evaluation

IF fine-tuning started from the representation-pretrained Xenium coordinate model, including its shared scale-aware transformer, central patch readout, and fine CNN. Villi were partitioned into training, validation, and evaluation sets within the two imaged sections. Training cells whose crops overlapped validation crops were excluded, leaving 5,822 training cells and 2,312 validation cells. Evaluation used 4,843 cells from 19 villi. Local, context, and fine crops were spatially disjoint between all three partitions.

IF adaptation first trained the fusion layers and coordinate heads for seven epochs at learning rate 3 × 10^-4^, with the shared ViT and fine CNN fixed. The validation-selected head-only model initialized four epochs of full-model training at 2 × 10^-5^. Minimum validation loss selected epoch 2 of head-only training and epoch 2 of full-model training, shown at epoch 9 on the cumulative training axis. Both stages used AdamW with weight decay 10^-4^, batch size 64, and gradient-norm clipping at 1. Intensity augmentation used probability 0.9, contrast 0.6–1.5, brightness shift 0, and blur sigma 0.1–2.5 pixels.

The Xenium model before IF adaptation and the IF-fine-tuned model were compared on identical crops from the 4,843 test cells, using the same training-derived reference scale. Spearman correlation was calculated for each model (**Extended Data Figure 4b**).

### Whole-tissue IF prediction and Immune Allocation Maps

IF predictions covered all 300,429 segmented cells: 163,992 DMSO and 136,437 RARi cells. QuPath classifications identified 9,104 DMSO and 1,994 RARi P14 cells; the remaining 289,331 cells formed the non-P14 reference population.

IMAPs placed predicted epithelial distance on the x-axis and predicted crypt-villus coordinate on the y-axis. Five fixed gates were applied: top intraepithelial, x ≤ 0.28 and y ≥ 0.40; top lamina propria, x > 0.28 and y ≥ 0.40; crypt intraepithelial, x ≤ 0.28 and y < 0.40; crypt lamina propria, x > 0.28 and 0.25 ≤ y < 0.40; and muscularis, x > 0.28 and y < 0.25. Gate names describe anatomical interpretations of the coordinate ranges. The same boundaries were used for P14 and non-P14 cells in both conditions.

For anatomical comparison, gate assignments were projected onto registered DAPI and E-cadherin images from both conditions (**Extended Data Figure 4d**). Fields extended the tissue magnifications to include underlying tissue (**Figure 4d**). E-cadherin was acquired in the Cy5 channel; display limits for each stain were shared across conditions. Each field was displayed with and without gate-colored cell centroids. Gate thresholds and memberships were identical to those used for IMAPs. The overlays provided a qualitative anatomical comparison.

Density was estimated on a 220 × 220 grid spanning [0, 1]^2^, smoothed with a Gaussian of sigma 3 bins and reflecting boundaries, then divided by population size and bin area. All four maps used color limits from zero to the largest grid density and contour levels at 0.1, 0.2, 0.4, 0.6, and 0.8 times that maximum.

Gate-specific abundance was expressed as P14 cells per 10,000 non-P14 cells in the same coordinate gate and condition. Raw P14 counts were also reported. Composition was the fraction of P14 cells assigned to each gate within a condition; differences were expressed as DMSO minus RARi percentage points. Comparisons were descriptive, with one section per condition.

### Metrics and reproducibility

Reference–prediction pairs with finite values were evaluated by Pearson and Spearman correlation, MAE, and the R^2^. Pearson r, MAE, and R^2^ were displayed for the coordinate predictions (**Figures 2d, 3c, and 4c**). Classifiers were evaluated by AUROC and average precision. Balanced accuracy, overall accuracy, precision, recall, and F1 were calculated at the stated thresholds.

## Acknowledgements

We thank Giovanni Galletti for providing immunofluorescence images for initial testing of SpatialTRACE-Image. Microscopy imaging was performed at the UCSD SOM Microscopy Core (NS047101, OD030505, OD036455). This research was supported by the Allen Institute, founded by Jody Allen – chair and co-founder of Allen Family Philanthropies, and the late Paul G. Allen – investor, philanthropist, and co-founder of Microsoft. We gratefully acknowledge their vision and generosity, which make this work possible.

## Materials and correspondence

Correspondence and requests for materials should be addressed to Maximilian Heeg.

## Author contributions

Conceptualization: A.M., A.W.G., and M.H.

Methodology: A.M., A.A.G., K.A., A.W.G., and M.H.

Investigation: A.M., A.A.G., K.A., and D.P.

Visualization: A.M., D.P., and M.H.

Funding acquisition: A.W.G. and M.H.

Project administration: A.M., A.W.G., and M.H.

Supervision: A.M., G.W.Y., A.W.G., and M.H.

Writing: A.M., D.P., A.A.G., K.A., G.W.Y., A.W.G., and M.H.

## Competing interests

A.W.G. is a co-founder of TCura Bioscience, Inc. G.W.Y. is a co-founder, member of the Board of Directors, Scientific Advisory Board member, equity holder and paid consultant for Eclipse BioInnovations. The other authors declare no competing interests.

## Data, code, and materials availability

Source code for the SpatialTRACE framework is publicly available on GitHub. Code for SpatialTRACE-Graph, including anatomical coordinate and region prediction from spatial transcriptomic data, is available at https://github.com/maximilian-heeg/SpatialTRACE-Graph. Code for SpatialTRACE-Image, including anatomical prediction from DAPI images, is available at https://github.com/amonell/SpatialTRACE-Image.

The spatial transcriptomics data analyzed in this study were previously generated and published in Reina-Campos et al., Nature (2025) and are available through the Gene Expression Omnibus (GEO) under accession GSE280895 (Xenium). Final code checkpoints, figure generation data, model weights, and the immunofluorescence images used for the retinoic acid receptor inhibitor analyses are deposited in 10.5281/zenodo.22850752. Additional data supporting the findings of this study are available from the corresponding authors upon reasonable request.

**Extended Data Figure 1.**
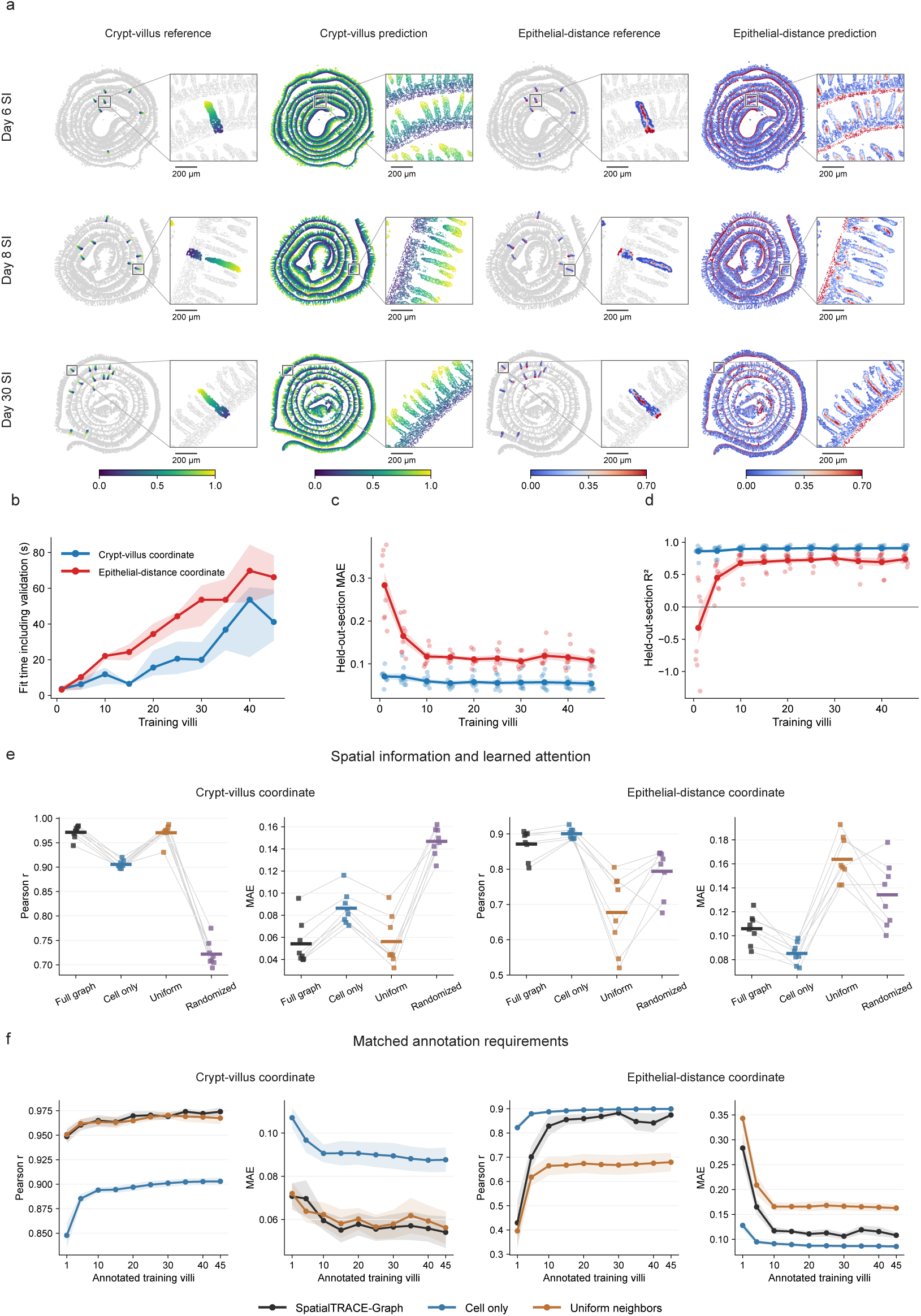
Sparse villus annotations support coordinate prediction across intestinal sections. **a,** Sparse references and dense graph predictions in small-intestine sections on days 6, 8, and 30 of lymphocytic choriomeningitis virus (LCMV) Armstrong infection. Rows show days 6, 8, and 30. Columns show crypt-villus references and predictions, followed by epithelial-distance references and predictions. Dense maps use models fitted to all sparse labels. **b–d,** Analyses use the eight section-disjoint folds and nested 1–45 training-villus subsets from Figure 2e. **b,** Model-fitting time for 160 fits, summarized by the median and interquartile range across folds. **c,** Mean absolute error (MAE) and **d,** coefficient of determination (R^2^). Points show held-out sections; lines and bands show mean ± standard error across eight sections. Villus counts refer to training annotations; validation uses an additional annotated section. Blue and red denote crypt-villus and epithelial-distance prediction, respectively. **e,** Full graph, cell-only, uniform-neighbor, and randomized-connection models trained separately using the same annotations, section splits, and validation-based selection. Randomization preserves the number of neighbors for every cell within its section. Points show held-out sections, gray lines connect matched sections, and colored bars show section means. **f,** Full graph, cell-only, and uniform-neighbor performance with matched 1–45 training-villus subsets and unchanged validation and test sections. Lines and bands show mean ± standard error across eight held-out sections. Panels **e** and **f** show Pearson correlation and MAE for both coordinates.

**Extended Data Figure 2.**
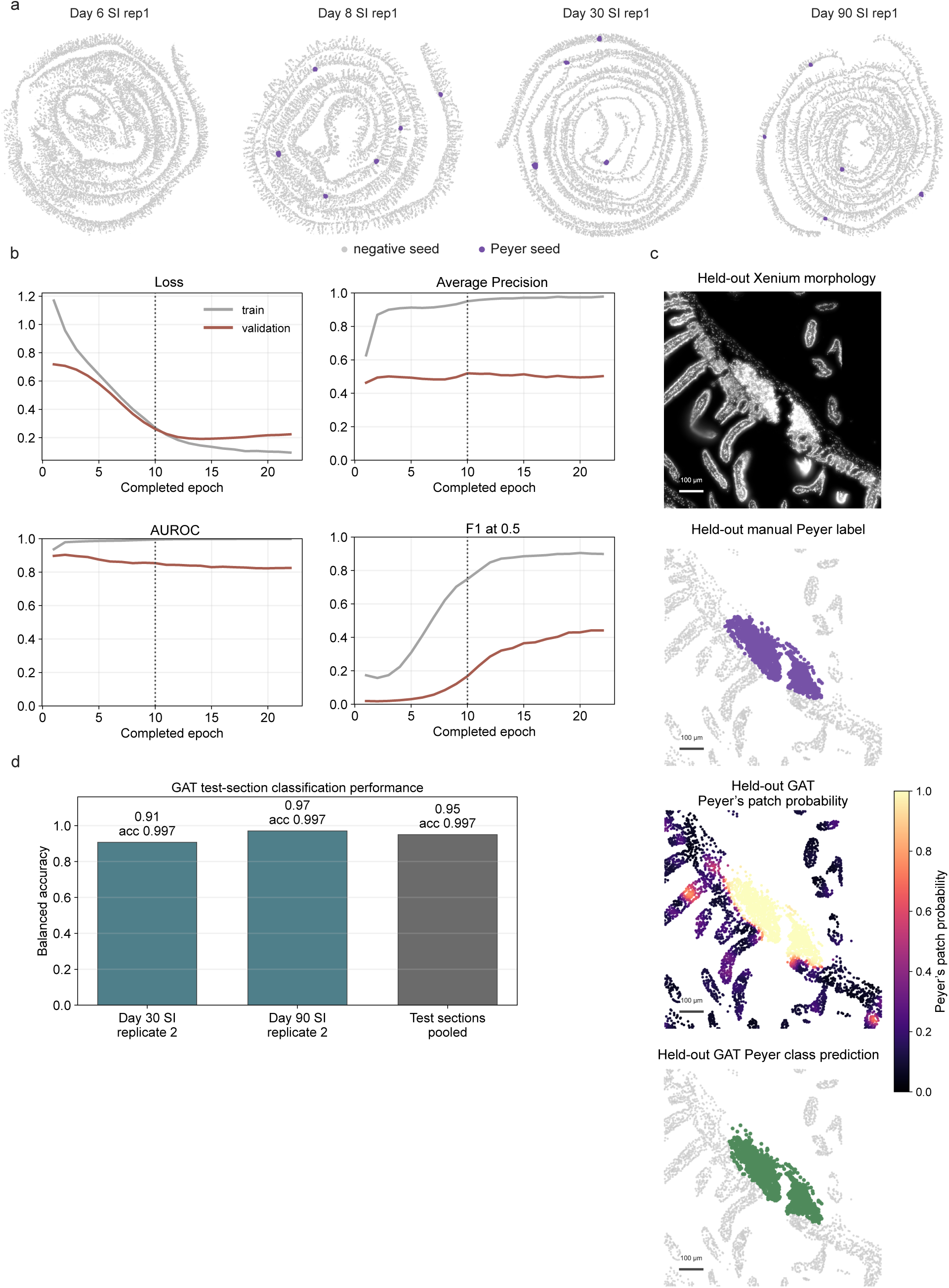
SpatialTRACE-Graph identifies discrete Peyer’s patch regions. **a,** Peyer’s patch training cells within annotated polygons (purple) and sampled cells outside the polygons (gray) in replicate-1 sections from days 6, 8, 30, and 90. Positive labels occur in the day 8, 30, and 90 sections. **b,** Training and validation loss, average precision, AUROC, and F1 at a probability threshold of 0.5 over 22 epochs. The dotted line marks epoch 10, selected by validation average precision. **c,** DAPI, manual labels, probabilities predicted by SpatialTRACE-Graph, and predicted classes in the day 90 replicate-2 test region. Classes use the threshold of 0.95 selected by validation F1. Scale bars, 100 µm. **d,** Balanced accuracy for each test section and the pooled 500,050 test cells. Supervised training, validation, and testing use separate sections.

**Extended Data Figure 3.**
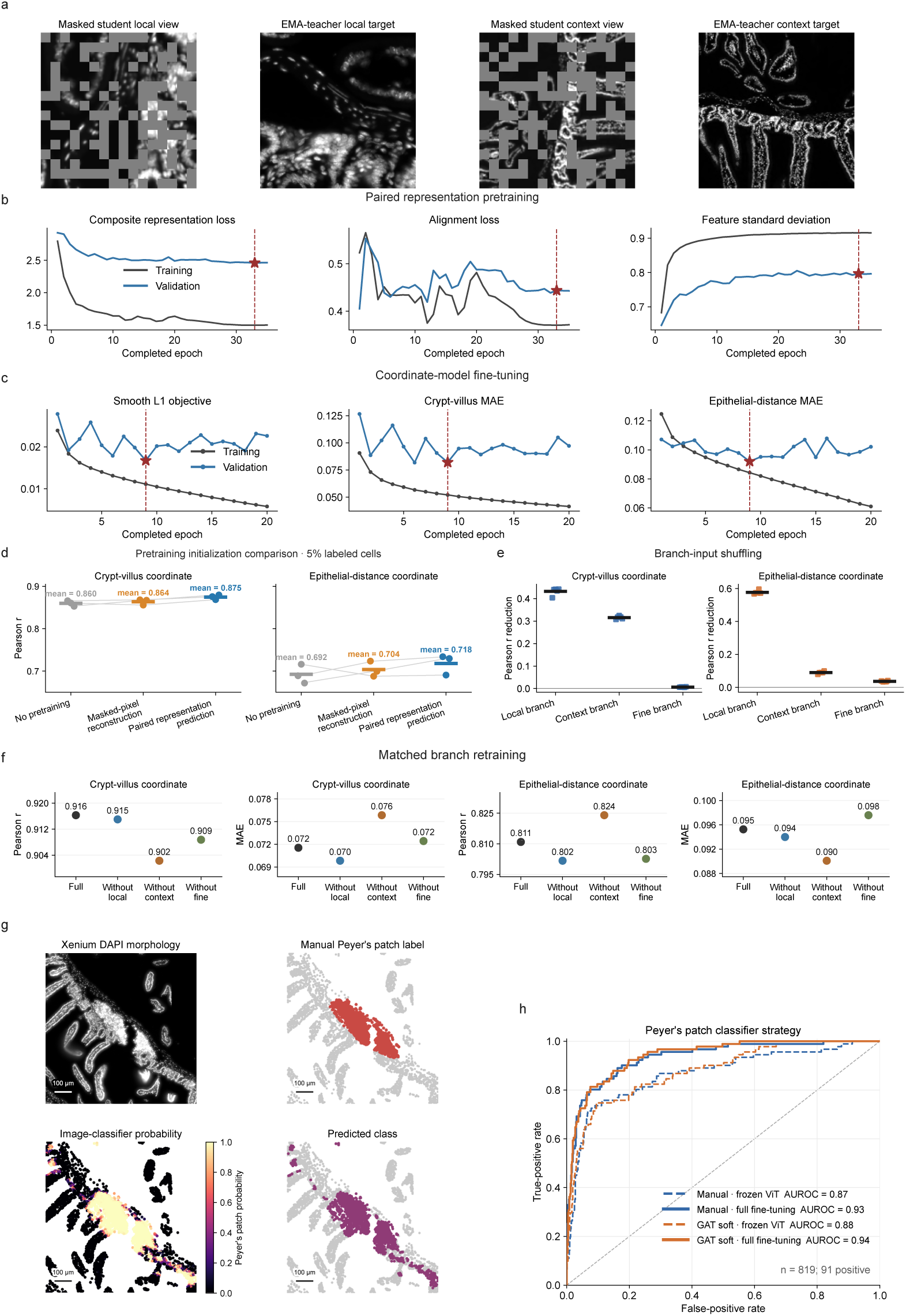
Pretraining, branch contributions, and image-based region classification. **a,** Paired student inputs with the same geometric and intensity transform at both scales, and clean teacher targets. Gray patches illustrate masking of approximately 40% of patch embeddings. **b,** Pretraining histories for total loss, alignment loss, and feature standard deviation. **c,** Fine-tuning history for the fully supervised coordinate model. Validation-selected models are marked in **b** and **c**. **d,** Comparison of random initialization, masked-pixel reconstruction, and paired representation prediction at 5% supervision. Architecture and fine-tuning splits are matched. Points show three fine-tuning runs, lines connect matched runs, and bars show means for the image-coordinate development set: 5,107 cells from four sections used for readout comparisons. **e,** Decrease in Pearson correlation after shuffling one crop type within evaluation batches while keeping the other inputs and model weights fixed. Points show five repeats; bars show means. This measures sensitivity to disrupted input pairing. **f,** Full model and models retrained without the local, context, or fine branch. Retained-branch initialization, training and validation cells, training settings, and checkpoint-selection criteria are matched. Points show Pearson correlation and MAE on the same image-coordinate development set as **d** and **e**, with one matched training run per architecture. **g,** DAPI, manual Peyer’s patch labels, image-classifier probabilities, and binary calls in the day 90 replicate-2 window. Calls use a probability threshold of 0.5. Scale bars, 100 µm. **h,** ROC curves compare manual and graph-derived soft targets with a frozen vision transformer (ViT) or full-model supervised fine-tuning.

**Extended Data Figure 4.**
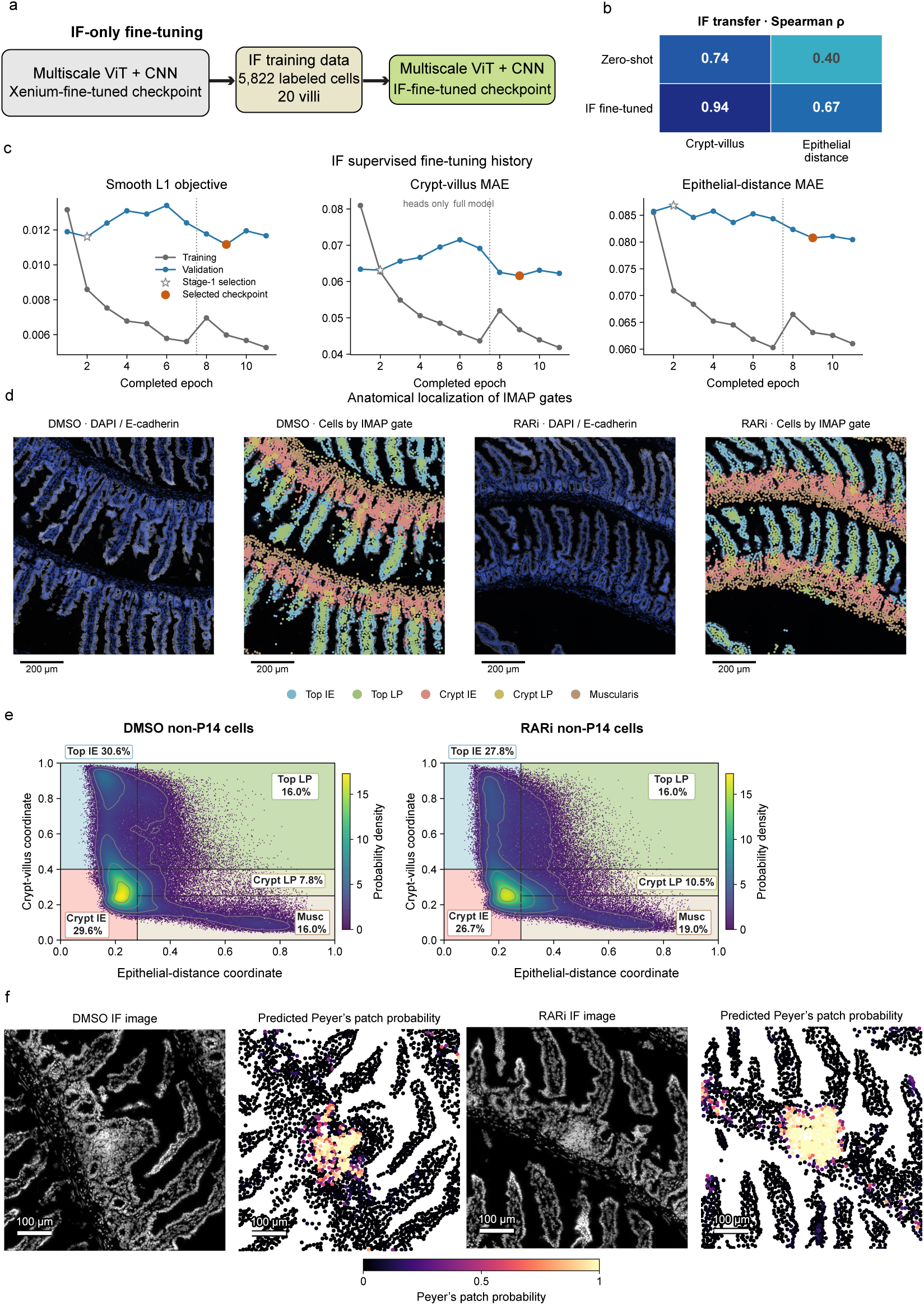
IF fine-tuning and anatomical mapping with SpatialTRACE-Image. **a,** Adaptation of the representation-pretrained Xenium coordinate model using 5,822 immunofluorescence (IF) training cells with spatially disjoint training, validation, and test crops. **b,** Spearman correlation before (Zero-shot) and after IF fine-tuning on the 4,843 test cells from Figure 4c, using identical crops and a common reference scale. **c,** Training and validation curves during head-only fine-tuning (epochs 1–7) and full-model fine-tuning (epochs 8–11). The star marks the head-only model selected at epoch 2 to initialize full-model fine-tuning. The orange point marks the final IF model selected at epoch 9. Each model was selected by the lowest validation loss within its training stage. **d,** DAPI (blue) and E-cadherin (gray; Cy5) images from DMSO and RARi sections beside the same fields with cells colored by Top IE, Top LP, Crypt IE, Crypt LP, or muscularis gate membership. IE, intraepithelial; LP, lamina propria. Gate assignments and boundaries match the IMAPs. Scale bars, 200 µm. **e,** Non-P14 IMAPs for DMSO (154,888 cells) and RARi (134,443 cells). Colors and contours show population-normalized probability density. Gates, density estimator, color limits, and contour levels match Figure 4f. Gate percentages use each condition’s non-P14 population as the denominator. Musc, muscularis. **f,** DAPI windows and Peyer’s patch probabilities in DMSO and RARi. The representation-pretrained image classifier described in **Extended Data Figure 3h** produces these scores. Both maps use the same linear 0–1 probability scale. Scale bars, 100 µm.

## Notes

https://github.com/maximilian-heeg/SpatialTRACE-Graph

https://github.com/amonell/SpatialTRACE-Image

https://zenodo.org/records/22850752

